# Recombination and selection against introgressed DNA

**DOI:** 10.1101/846147

**Authors:** Carl Veller, Nathaniel B. Edelman, Pavitra Muralidhar, Martin A. Nowak

## Abstract

DNA introgressed from one species into another is typically deleterious at many genomic loci in the recipient species. It is therefore purged by selection over time. Here, we use mathematical modeling and whole-genome simulations to study the influence of recombination on the purging of introgressed DNA. We find that aggregate recombination controls the genome-wide rate of purging in the first few generations after admixture, when purging is most rapid. Aggregate recombination is quantified by 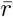, the average recombination rate across all locus pairs, and analogous metrics. It is influenced by the number of crossovers (i.e., the map length) and their locations along chromosomes, and by the number of chromosomes and heterogeneity in their size. A comparative prediction of our analysis is that species with fewer chromosomes should purge introgressed DNA more profoundly, and therefore should exhibit a weaker genomic signal of historical introgression. With regard to patterns across the genome, we show that, in heterogametic species with autosomal recombination in both sexes, more purging is expected on sex chromosomes than on autosomes, all else equal. The opposite prediction holds for species without autosomal recombination in the heterogametic sex. Finally, we show that positive genomic correlations between local recombination rate and introgressed ancestry, as recently observed in several taxa, are likely driven not by recombination’s effect in unlinking neutral from deleterious introgressed alleles, but rather by its effect on the rate of purging of the deleterious alleles themselves.

**Note on this version:** An earlier version of this manuscript had two parts: (1) Calculations of the variance of genetic relatedness between individuals with particular pedigree relationships, taking into account the randomness of recombination and segregation in their pedigree. (2) An investigation of the rate of purging of introgressed DNA following admixture, based in part on results from part (1). Part (1) has since been published as Veller et al. (2020). The present manuscript has been reconfigured to focus on part (2).

## 1 Introduction

It has become clear in recent years that hybridization and subsequent genetic introgression are common features of the evolutionary histories of many species, including our own (Taylor and Larson 2019; Edelman and Mallet 2021). It has therefore become a major focus of evolutionary genetics to understand the impact of introgression on the population genomics of species, and conversely to learn about historical admixture from genomic patterns of introgressed ancestry (Moran et al. 2021).

Introgressed DNA is typically deleterious in the recipient species. This can be for a number of reasons: donor-species alleles could be maladapted to the recipient species’ ecology (Schluter 2009) or genome (Dobzhansky 1937; Muller 1942; Orr 1995), and the donor species could carry a higher genetic load than the recipient species (Juric et al. 2016; Harris and Nielsen 2016). Importantly, the deleterious effect of introgressed ancestry is often spread across a large number of loci. For example, it has been estimated that Neanderthal alleles were deleterious in humans at ~1,000 loci (Juric et al. 2016; Harris and Nielsen 2016); comparable estimates have been obtained for other species [e.g., Schumer et al. (2014); Aeschbacher et al. (2017)].

Since introgressed alleles initially appear in the recipient population in perfect linkage disequilibrium, recombination will clearly be influential in determining their fate. The influence of recombination on the purging of deleterious introgressed alleles can be understood in two complementary ways. The first is that recombination breaks up the initial long blocks of introgressed DNA, which are strongly selected against, into smaller and smaller blocks, which are more weakly selected against (Barton 1983). The second is that recombination, by distributing introgressed ancestry more and more evenly among more and more individuals, reduces ancestry variance over time and thus reduces the efficiency of selection against deleterious introgressed ancestry (Harris and Nielsen 2016). Both views reveal that recombination should, over time, reduce the rate at which deleterious introgressed ancestry is purged.

This process has been well studied in the context of stylized genetic maps (Bengtsson 1985) and small genomic segments [e.g., Barton (1983); Barton and Bengtsson (1986)]. However, in light of recent evidence that introgressed alleles can be deleterious at many loci throughout the genome, and that many species—with a great diversity of recombination processes—have experienced introgression, it is worthwhile revisiting the role that recombination plays in the purging of introgressed DNA, taking a genome-wide perspective.

Here, we use mathematical models and computer simulations to study the role of recombination in selection against introgressed DNA. First, we ask what features of the recombination process affect the efficiency with which introgressed ancestry is purged genome-wide. Cross-species variation in these features would be expected to predict variation in the retention of introgressed ancestry, and thus the strength of the genomic signal of historical admixture. Second, inspired by recent empirical findings in several taxa that introgressed ancestry is preferentially retained in high-recombination regions of the genome (Brandvain et al. 2014; Schumer et al. 2018; Martin et al. 2019; Edelman et al. 2019; Calfee et al. 2021), we investigate the mechanisms that generate these within-genome correlations between recombination rate and introgressed ancestry.

## 2 Model

We study a model in which selection acts additively against introgressed DNA: if a proportion *p* of an individual’s genome is introgressed, its relative fitness is 1 − *pS*. This is the additive version of the model in Barton (1983), and corresponds to a situation where introgressed alleles are deleterious in the recipient species at a large number of loci, with fitness effects additive within and across loci. The loci at which introgressed alleles are deleterious are assumed to be spaced uniformly along the physical map of the genome, although this assumption can be relaxed with minimal change to the interpretation of our results.

Notice that, in assuming that an individual’s fitness is determined by the fraction of its genome that is introgressed, we have ignored the distinction between deleterious and neutral (or beneficial) introgressed alleles. If introgressed alleles are deleterious at sufficiently many loci throughout the genome, this assumption will have little effect on the overall rate at which introgressed ancestry is purged (Fig. S5). To understand how and when recombination can cause differential retention of deleterious and neutral introgressed DNA, we later carry out simulations with neutral loci included amongst the deleterious loci.

Similar to previous work [e.g., Harris and Nielsen (2016); Juric et al. (2016); Steinrücken et al. (2018)], we assume that hybridization occurs as a pulse in a single generation, such that a certain fraction of individuals in the next generation are F1 hybrids, each with half its genome introgressed. Mating is assumed to be random with respect to ancestry. For simplicity, we ignore sex chromosomes at first. Finally, we assume that the population is large enough that drift can be ignored (in our simulations, the population size *N* = 100,000).

All simulations were carried out in SLiM 3 (Haller and Messer 2019).

## 3 Species differences in the genome-wide rate of purging

Figure 1 shows the genome-wide purging of introgressed DNA in simulations of the model described above, under the recombination processes of humans and *Drosophila melanogaster* [using linkage maps produced by Kong et al. (2010) and Comeron et al. (2012) respectively]. All other parameters are identical between the two cases, and were chosen to resemble parameters inferred for Neanderthal-human introgression (Harris and Nielsen 2016; Juric et al. 2016): the initial introgression fraction is 5%, introgressed alleles are deleterious at 1,000 loci, and F1 hybrids suffer a 20% fitness reduction. Simulations were run for 2,000 generations, approximately the number of human generations since Neanderthal introgression. Several features of the trajectories in Figure 1 are noteworthy.

**Figure 1:**
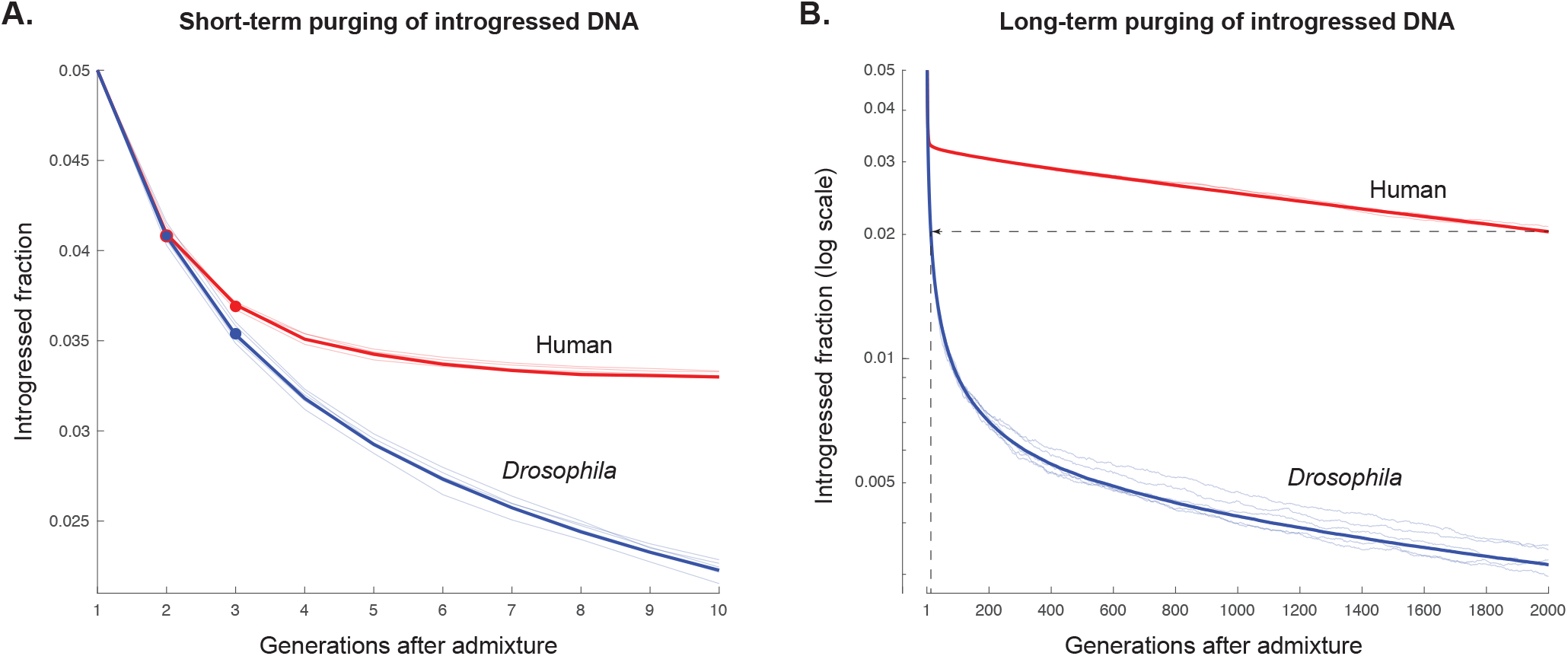
Genome-wide purging of introgressed DNA following an admixture pulse, under the recombination processes of humans and *Drosophila melanogaster*. The recombination process of *D. melanogaster* causes much more introgressed ancestry to be purged in the first few generations after admixture (**A**), and thus by later generations as well (**B**), because it is associated with a lower aggregate recombination rate, driven predominantly by the small karyotype of *D. melanogaster* (2 major autosomes) relative to humans (22 autosomes). The dots in **A** are analytical predictions from Eqs. (6) and (10). The bold lines are trajectories averaged across 100 replicate simulations; the faint lines are representative trajectories. The dotted line in **B** marks that *Drosophila* purges as much introgressed DNA in 12 generations as humans do in 2,000 generations.

First, in both humans and *Drosophila*, introgressed DNA is purged very rapidly in the first few generations after admixture. The rate of purging subsequently decreases. This effect, predicted by previous calculations (Bengtsson 1985) and observed in recent simulations (Harris and Nielsen 2016; Petr et al. 2019), is so extreme that, for both species, most of the purging that has eventually occurred after 2,000 generations actually occurred in the first 5 generations.

Second, *Drosophila* purges introgressed DNA much more efficiently than humans. After 2,000 generations, *Drosophila* has purged 94% of the initial introgressed fraction, while humans have purged only 59%. Put differently, the amount of introgressed DNA that it takes humans 2,000 generations to purge, *Drosophila* has purged after just 12 generations. Therefore, the recombination process of *D. melanogaster* is a much more effective barrier to gene flow than the recombination process of humans.

The observations above are robust to alternative specifications of our model in which the fitness effects of introgressed alleles are variable across loci (Fig. S3) or driven by Dobzhansky-Muller incompatibilities (Fig. S4).

Our aim in the following subsections is to derive analytical expressions that help us to understand these observations.

### 3.1 Preliminary calculations

Let *Z_t_* be the fraction of a random generation-*t* zygote’s genome that is introgressed, and let *G_t_* be the introgressed fraction of a random gamete produced by generation-*t* adults. Let *A_t_* be the proportion of generation-*t* zygotes that have hybrid ancestry (in the pedigree sense—i.e., some of them might nevertheless have inherited no introgressed DNA), and *A′_t_* the corresponding proportion of adults after viability selection (and therefore also the proportion of successful gametes produced by generation *t*). Let 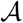 signify the property of having hybrid ancestry (which applies to a proportion *A_t_* of zygotes and *A′_t_* of adults and gametes). A graphical representation of the variables used in these calculations is given in Fig. S2.

We are interested in how the overall fraction of introgressed DNA, 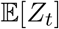, is reduced over time by selection—this is the process displayed in Fig. 1. We show in SI Section S1 that the fraction of introgressed DNA purged in the *t*-th generation is

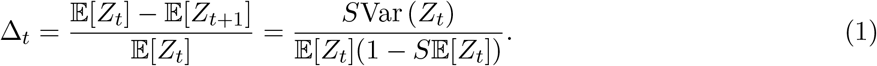

Therefore, to understand what factors govern the rate of purging of introgressed DNA, we must understand what factors govern the trajectory of Var(*Z_t_*), the variance across individuals in how much introgressed DNA they carry (Harris and Nielsen 2016). Var(*Z_t_*) can be decomposed into two components: a contribution from the fact that some individuals have introgressed ancestry and some do not, and a contribution from variance among those individuals that do have introgressed ancestry. We shall call these the ‘between group’ and ‘within group’ components of the overall variance, respectively. The precise decomposition is

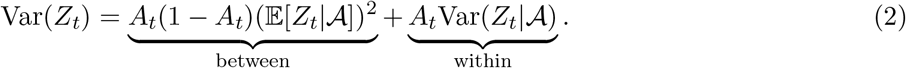

An analogous decomposition holds for genetic ancestry variance across gametes:

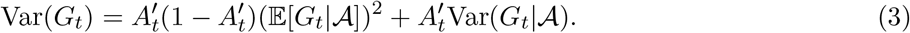

We begin by calculating Var(*Z*_1_) and Var(*Z*_2_). For later *t*, we provide an approximate calculation of Var(*Z_t_*) in the special case where selection is weak and the initial introgressed fraction is small.

### 3.2 Short-term purging of introgressed DNA

#### Selection in the first generation

A zygote in the first generation after admixture either carries no introgressed DNA (probability 1− *A*_1_) or is an F1 hybrid, with half of its genome introgressed (probability *A*_1_). Therefore,

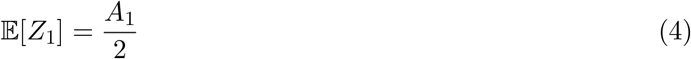

and

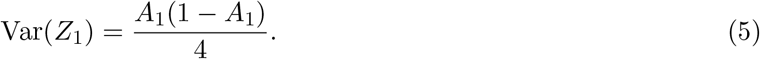

Notice that all of the ancestry variance in generation 1 is due to differences between hybrids and non-hybrids—there is no ‘within group’ contribution, because all hybrids have the same introgressed fraction (1/2).

From Eq. (1), the proportion of introgressed DNA removed by selection in the first generation is

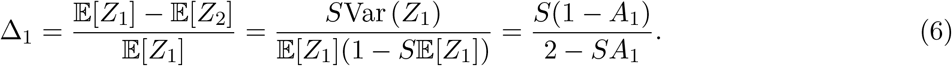

This expression is, of course, independent of the recombination process, explaining why the introgressed fraction in generation-2 zygotes in our simulations is the same for humans and *Drosophila* (Figure 1A).

#### Selection in the second generation

Some generation-2 zygotes have F1 hybrid parents; recombination in these parents will generate ancestry variance among their offspring. From Eq. (1) in Veller et al. (2020), among those gametes produced by generation-1 hybrids,

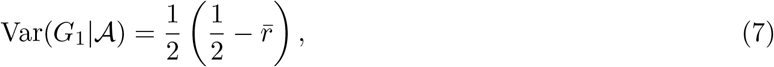

where 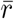 is the average recombination rate across all pairs of loci in the genome (Veller et al. 2019). When the recombination process differs between the sexes—as is usually the case (Lenormand and Dutheil 2005; Sardell and Kirkpatrick 2020)—the sex-averaged value of 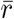 applies in Eq. (7).

Noting that 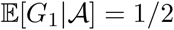, and applying Eq. (3), we find that

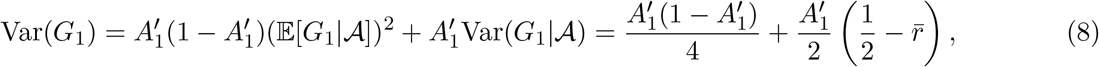

where 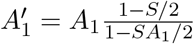 is the fraction of generation-1 gametes that derive from hybrid parents.

Since a zygote’s introgressed fraction is the average of the introgressed fractions of the gametes that produced it, and since we have assumed that mating is random with respect to ancestry,

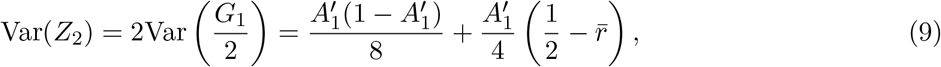

where the first term is the ‘between group’ contribution and the second term is the ‘within group’ contribution.

Finally, we observe that 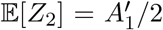, and substitute Eq. (9) into Eq. (1) to find the proportion of introgressed DNA that is purged in the second generation after admixture:

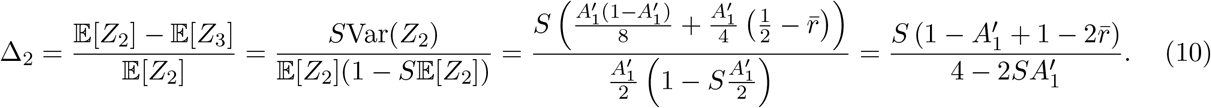

Eq. (10) reveals that the rate of purging of introgressed DNA in the second generation after admixture depends on the aggregate recombination rate, quantified by 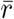. 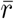 can be measured from various kinds of data, including cytological data of crossover positions at meiosis I, sequence data from gametes, and linkage maps (Veller et al. 2019). 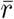 is influenced by several features of the recombination process: the number of chromosomes and heterogeneity in their size, the number of crossovers and their locations along the chromosomes, and the spatial relationships among crossovers (i.e., crossover interference). In most species, the dominant contribution to 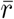 is from independent assortment of chromosomes at meiosis (Crow 1988; Veller et al. 2019). Therefore, the primary cause of variation in 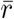 aross taxa is variation in chromosome number, with crossovers playing a secondary role.

These considerations explain a key feature of the trajectories in Figure 1A—that in the first few generations after admixture, *D. melanogaster* purges introgressed DNA much more rapidly than humans do. *D. melanogaster* has only two major autosomes, the independent assortment of which at meiosis contributes relatively little genetic shuffling. Furthermore, crossing over is absent in males. In Veller et al. (2020), we estimated a sex-averaged autosomal value of 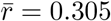 for *D. melanogaster*, substantially less than the theoretical maximum of 1*/*2. In contrast, humans have 22 autosomes, the independent assortment of which generates much genetic shuffling. Veller et al. (2019) estimated a sex-averaged autosomal value of 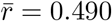 in humans, close to the theoretical maximum. Substituting these values into Eq. (10), we obtain predictions of how much introgressed DNA is retained by humans and by *Drosophila* in the third generation after admixture. These analytical predictions agree well with our simulations (Figure 1A).

We conclude that, relative to humans, *Drosophila* purges introgressed DNA more efficiently in the early generations after admixture because its aggregate recombination process generates substantially less genetic shuffling, owing primarily to its small karyotype and the absence of crossing over in males. Generalizing, this suggests that karyotypic variation across species could be a primary driver of species differences in the retention of introgressed DNA and thus the strength of the genomic signal of historical introgression (Fig. S6).

### 3.3 Long-term purging of introgressed DNA

After the third generation post-admixture, the complex interaction of recombination and selection prevents tractable analytical calculations in the general case. Nevertheless, we can make analytical progress in understanding the impact of recombination on the purging of introgressed DNA in these later generations by considering a special case where the initial introgressed fraction is small (*A*_0_ ≪ 1) and selection against introgressed ancestry is weak (*S* ≪ 1).

In this case, selection does not appreciably limit the number of descendants of the initial hybrids, so that the fraction of the population with introgressed ancestry grows exponentially:

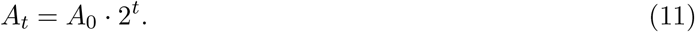

Concomitantly, because selection does not appreciably deplete the overall fraction of introgressed DNA 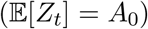, the average fraction among those individuals with introgressed ancestry declines exponen-tially:

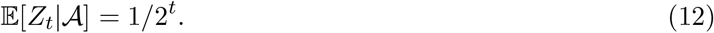

Each generation-*t* individual with introgressed ancestry descends from a single introgressing ancestor in generation 0, and, because selection has not appreciably affected the distribution of genetic ancestry among these generation-*t* individuals, the ancestry variance among them is equal to the variance of a random individual’s genetic relatedness to one of its *t*-th degree great-grandparents. This quantity is calculated in Veller et al. (2020):

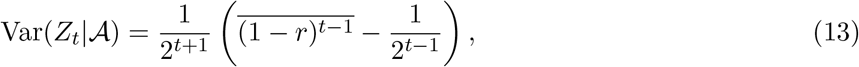

where 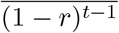 is the average value of (1 − *r*_*ij*_)^*t*−1^ taken over all locus pairs (*i, j*), with *r_ij_* the sex-averaged recombination rate between loci *i* and *j* [notice that this expression is not the same as 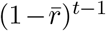]. Substituting Eq. (13) into Eq. (2), we find that the overall variance of genetic ancestry in generation *t* is

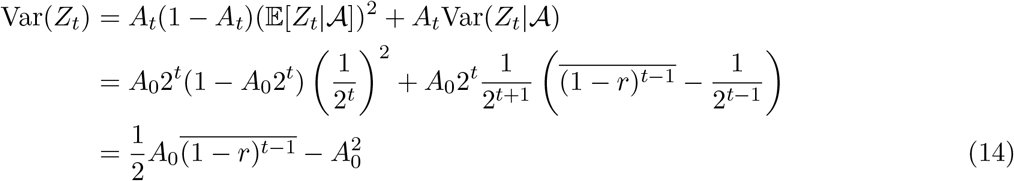

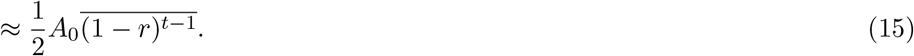

Substituting this result into Eq. (1), we find that the rate of purging of introgressed DNA in generation *t* is

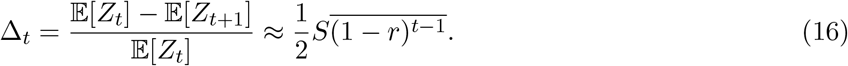

We can make several observations from Eqs. (15) and (16). First, because the terms (1 − *r*_*ij*_)^*t*−1^ that contribute to the average 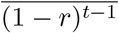 decline exponentially over time (since 1 − *r_ij_* < 1), the variance of introgressed ancestry declines over time [Eq. (15)], and therefore so does the rate of purging of introgressed DNA [Eq. (16); Fig. 1].

Second, Eq. (16) is informative of the genomic scales of recombination that most influence the rate of purging at different timepoints after admixture. Consider again the terms (1 − *r*_*ij*_)^*t*−1^ that contribute to 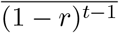. Some of these terms come from pairs of loci on different chromosomes (*r_ij_* = 1*/*2), while others come from pairs of loci linked along the same chromosome (*r_ij_* < 1*/*2). The term (1 − *r*_*ij*_)^*t*−1^ from each unlinked locus pair is smaller than the term from each linked locus pair, but, in most species, there are many more unlinked locus pairs than linked locus pairs (Crow 1988; Veller et al. 2019). Therefore, it is ambiguous whether unlinked or linked locus pairs in total contribute more to 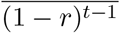 —that is, whether the rate of purging of introgressed DNA is influenced more by independent assortment of chromosomes or by crossing over.

In the early generations after admixture, when *t* is small, a term (1 − *r*_*ij*_)^*t*−1^ from an unlinked locus pair cannot be much smaller than a term from a linked pair of loci: when *t* = 2, for example, the value in the former case is 1/2 while the maximum possible value in the latter case is 1. Therefore, when the great majority of locus pairs are unlinked (as in most species), independent assortment will be the dominant contributor to 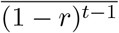 in the early generations after admixture. This is consistent with our main result in the previous subsection, and its associated prediction that variation in chromosome number is a major cause of species differences in the early rate of purging of introgressed DNA.

In later generations, as *t* increases, each term (1 − *r*_*ij*_)^*t*−1^ that contributes to 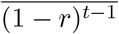 shrinks exponentially, with terms from unlinked loci (and loci genetically far apart) shrinking much more rapidly than terms from tightly linked loci. Thus, many generations after admixture, only terms from tightly linked loci contribute meaningfully to 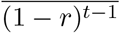 Therefore, in these later generations, species differences in the rate of purging are expected to be driven by fine-scale recombination rates between tightly linked locus pairs.

### 3.4 Intuition

Our main result above, that species with lower aggregate recombination rates purge introgressed DNA more efficiently, has a simple intuition. Initially, introgressed DNA appears in the recipient population in long linkage blocks, each the size of a haploid genome. Recombination influences the subsequent rate of purging of introgressed DNA because it breaks up these intial long blocks into smaller, variably-sized blocks. A species with a low aggregate recombination rate will maintain introgressed DNA in longer and/or more variably sized blocks, and will therefore purge introgressed DNA more efficiently (Barton 1983).

The average size of the blocks into which recombination chops the initial long blocks is determined predominantly by the number of chromosomes and crossovers, while the variability of block size is determined largely by heterogeneity in the sizes of chromosomes and the spatial arrangement of crossovers along chromosomes. To see this, note that a crossover anywhere along a block of introgressed DNA will break that block up into two smaller blocks with an average size of half the parent block. The position of the crossover along the block, however, will determine how variably sized the two descendant blocks are. Thus, all else equal, a species with more chromosomes and/or more crossovers will purge introgressed DNA less efficiently, because it more rapidly breaks up the initial introgressed blocks into smaller blocks, which are less deleterious. And, all else equal, a species that situates crossovers near the tips of chromosomes will purge introgressed DNA more efficiently than a species with a uniform distribution of crossovers, because, in the former species, the resulting blocks of introgressed DNA will vary greatly in size.

In SI Section S2, we analyze a simple model that distinguishes the contributions of these two factors—the mean and variance of introgressed block length—to ancestry variance across individuals and thus the rate at which introgressed DNA is purged. The model applies best when sufficiently many generations have elapsed since the initial admixture pulse that different blocks of introgressed DNA can be assumed to have been inherited independently. Applying this model to the recombination process of *D. melanogaster*, we find that (i) the model accurately predicts the rate of purging of introgressed DNA (Fig. S1A), (ii) the contribution of block length variance is greatest in the early generations after admixture, when crossover locations and variation in chromosome size have greatest effect (Fig. S1B), and (iii) block number variance across individuals—which in this model is proportional to the average block length—becomes the most important contributor to ancestry variance in the long term (Fig. S1B). These results are again consistent with the view that aggregate recombination—as quantified by 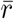 and analogs—determines the short-run rate of purging of introgressed DNA, while fine-scale recombination rates determine the long-run rate of purging.

## 4 Variation in the rate of purging across the genome

The intuition above applies not only to different species, but also to different regions within a given species’ genome. In genomic regions with low aggregate rates of recombination, deleterious introgressed alleles will be maintained in larger and/or more variably sized blocks, and will therefore be purged more efficiently. Thus, for example, a chromosome that typically receives few crossovers per meiosis will, all else equal, purge introgressed DNA more rapidly than a chromosome that receives more crossovers (Fig. S7A). Similarly, if two chromosomes receive the same number of crossovers per meiosis, but crossovers tend to be terminally situated on the one chromosome and uniformly distributed along the other, then the chromosome with the more terminal distribution will purge introgressed DNA more rapidly, owing to its lower chromosomespecific value of 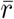 (Fig. S7B).

Below, we study two particularly interesting implications of recombination differences across the genome for the purging of introgressed DNA.

### 4.1 Sex chromosomes

Sex chromosomes often show unusual genetic signatures of admixture. Most notably, X and Z chromosomes tend to retain less introgressed ancestry than autosomes (Martin and Jiggins 2017). Several factors could account for the lower signal of admixture on X/Z chromosomes, including their enrichment for alleles that reduce hybrid fitness (Charlesworth et al. 1987; Presgraves 2008) and their hemizygous expression in one sex (Turelli and Orr 1995). Here, we explore an additional factor that affects the retention of introgressed ancestry on sex chromosomes versus autosomes: recombination differences.

In species with a degenerate sex-specific chromosome (the Y or W), recombination along most of the X/ Z chromosome is typically restricted to the homogametic sex (XX females or ZZ males). This affects the average rate at which the X/Z chromosome recombines, relative to the autosomes. First, consider a species with autosomal recombination in both sexes (i.e., most heterogametic species). Unless the X/Z chromosome recombines at a substantially elevated rate in the homogametic sex, its lack of recombination in the heterogametic sex will ensure that, on average across generations, it experiences less recombination than the autosomes. Now consider a species without autosomal recombination in the heterogametic sex (e.g., *Drosophila*, Lepidoptera). Assuming an even sex ratio, in a given generation, one half of the copies of each autosome are present in the homogametic sex and therefore recombine. In contrast, two thirds of the copies of the X/Z chromosome are present in the homogametic sex and therefore recombine. So, in this case, the X/Z chromosome actually experiences more recombination than the autosomes.

Therefore, all else equal, we expect sex chromosomes to retain less introgressed DNA than autosomes in species with autosomal recombination in the heterogametic sex; but in species without autosomal recombination in the heterogametic sex, we expect the sex chromosomes to retain more introgressed DNA than the autosomes. These predictions were confirmed in simulations of our model augmented to include sex chromosomes, for the recombination processes of humans (heterogametic sex does recombine) and *D. melanogaster* (heterogametic sex does not recombine), and for more stylized recombination processes as well (Fig. 2).

**Figure 2:**
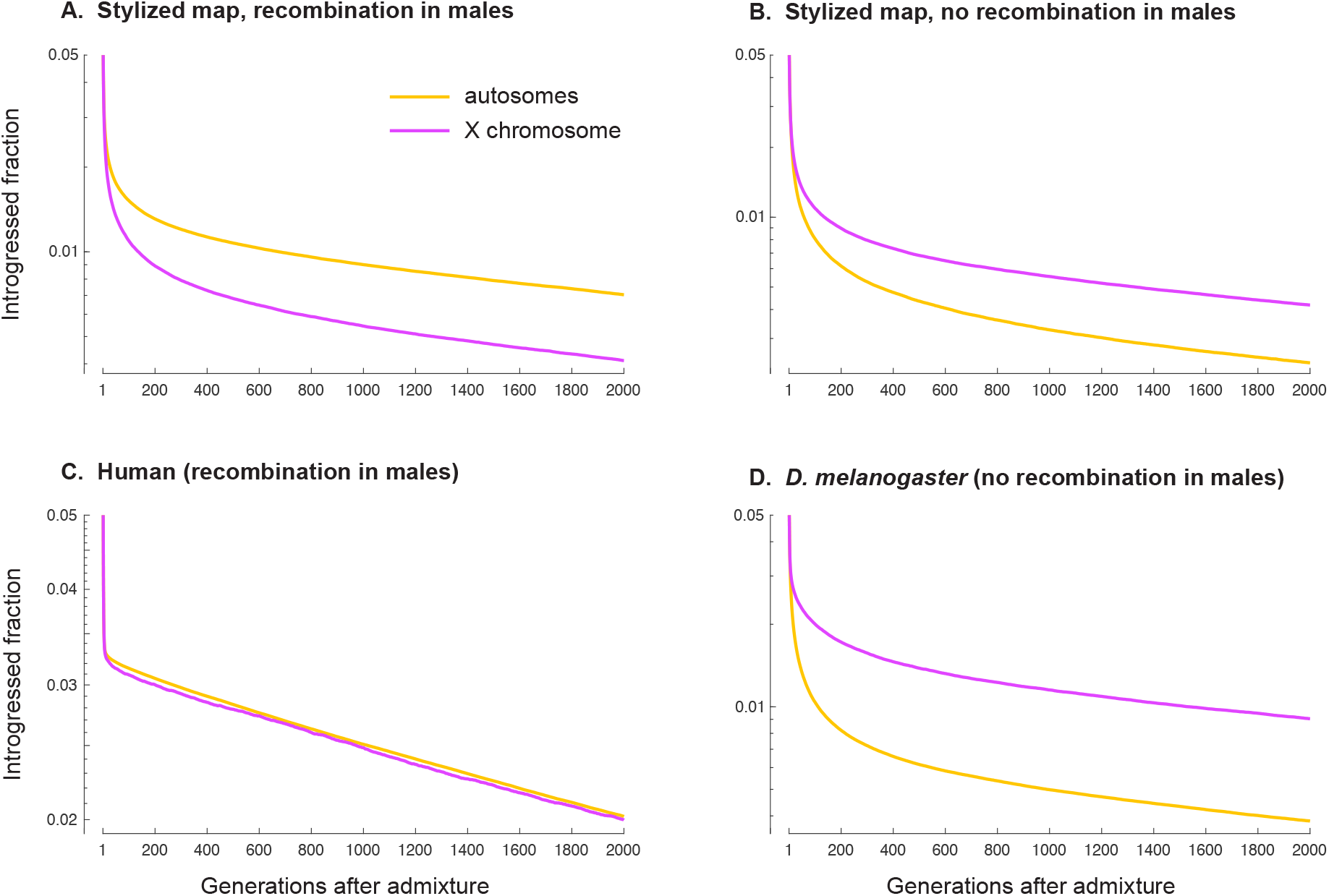
Recombination differences between sex chromosomes and autosomes generate differences in the rate of purging of introgressed DNA. **A**,**C**. In species with autosomal recombination in both sexes, the lack of recombination along the sex chromosome in the heterogametic sex (here, the X chromosome in males) leads to a lower average rate of recombination than on autosomes. The sex chromosome therefore purges introgressed DNA more rapidly than the autosomes, all else equal. **B**,**D**. In species without autosomal recombination in the heterogametic sex, the sex chromosome has a higher average rate of recombination than the sex chromosomes, because 2/3 of its copies are in the (recombining) homogametic sex, compared to only 1/2 of autosomes. Therefore, in such cases, all else equal, the sex chromosome purges less introgressed DNA than the autosomes. The simulations here assume full dosage compensation of the sex chromosome in the heterogametic sex, such that the relative fitness of a heterogametic individual with autosomal and sex-linked introgressed fractions *p_A_* and *p_X_* is 1 − *S*(*p_A_* + 2*p_X_*)*/*3. Trajectories are averages across 100 replicate simulations. The stylized genomes in **A** and **B** involve one autosome, of equal size to the sex chromosome (500 deleterious loci each), with the sex chromosome and the autosome each receiving, on average, two crossovers in the homogametic sex, with the location of each crossover sampled independently and uniformly along its chromosome. In **A**, the autosomal recombination process is identical in the heterogametic and homogametic sexes.

Thus, recombination differences alone can generate differences in the retention of introgressed DNA between sex chromosomes and autosomes. Of course, recombination differences cannot fully explain the ancestry differences actually observed between sex chromosomes and autosomes, since additional features of sex chromosomes are known to be important in this regard (Coyne and Orr 2004; Payseur et al. 2018). Indeed, in some cases, the predicted impact of recombination differences is opposite in direction to the observed ancestry disparity between sex chromosomes and autosomes. For example, in *Heliconius* butterflies, with female heterogamety and no autosomal recombination in females, Z chromosomes are especially depleted for introgressed ancestry [Van Belleghem et al. (2018); Martin et al. (2019); though see Zhang et al. (2016) for an interesting exception]. This is in spite of the Z having a higher average recombination rate than the autosomes, per the argument above. In such cases, the factors that underlie the reduced retention of introgressed ancestry on the sex chromosome must be even stronger than previously thought, since they must work against the countervailing effect of recombination differences between the sex chromosomes and the autosomes.

### 4.2 Correlations between regional recombination rate and introgressed ancestry

Several recent studies, encompassing a broad diversity of taxa, have identified a positive genomic correlation between local recombination rate and the amount of introgressed ancestry (Brandvain et al. 2014; Schumer et al. 2018; Martin et al. 2019; Edelman et al. 2019; Calfee et al. 2021). That is, regions of the genome that experience less recombination tend to retain less introgressed DNA. In trying to understand these correlations, it is important to note that even if introgressed alleles are deleterious at many loci throughout the genome, the overwhelming majority of introgressed DNA is expected to be neutral. Therefore, the positive correlations that have been reported are driven by heterogeneity across the genome in the retention of neutral introgressed alleles (Schumer et al. 2018).

For a neutral introgressed allele ultimately to survive in the recipient population, it must recombine away from its flanking deleterious introgressed alleles before those deleterious alleles are eliminated by selection (Bengtsson 1985; Barton and Bengtsson 1986). Recombination affects this process in two ways (Schumer et al. 2018): (i) it affects the rate at which deleterious introgressed alleles are purged, and (ii) it affects the rate at which neutral introgressed alleles dissociate from their flanking deleterious alleles. These two effects of recombination constitute two distinct mechanisms by which local recombination rates across the genome come to be positively correlated with local ancestry. (To see that the two mechanisms are conceptually distinct, consider the case where every neutral introgressed allele is completely linked to a deleterious introgressed allele. Then no neutral allele ever dissociates from its flanking deleterious alleles, so that, eventually, all neutral alleles will be purged. Despite this lack of unlinking, in the long period before purging is completed, there will be a positive correlation between recombination rate and neutral introgressed ancestry, because a greater fraction of deleterious ancestry [and linked neutral ancestry] will have been purged in regions of low recombination.) The mechanisms are concordant: In regions of low recombination, deleterious alleles are purged more rapidly, and neutral alleles remain linked to their flanking deleterious alleles for a longer time.

The relative contributions of these two mechanisms to the overall correlation between recombination rate and introgressed ancestry can be quantitatively distinguished using the following decomposition, a graphical representation of which is given in Fig. S8:

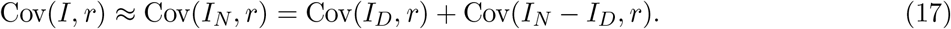

Here, *I* is the overall introgressed fraction in a given genomic window, *I_N_* and *I_D_* are the introgressed fractions at neutral and deleterious loci respectively, and *r* is some measure of the recombination rate within the window. The first term on the right hand side of Eq. (17), Cov(*I_D_, r*), captures the direct effect of recombination on the rate of purging of deleterious introgressed alleles—effect (i) above. The second term, Cov(*I_N_* − *I_D_, r*), captures the effect of recombination in unlinking neutral from deleterious introgressed alleles, thus decoupling their frequency trajectories—effect (ii) above. The relative importance of the ‘unlinking effect’ is given by Cov(*I_N_* − *I_D_, r*)*/*Cov(*I, r*).

To analyze the genomic correlation between recombination rate and introgressed ancestry in our simulations, we augmented our model to include neutral loci between the loci at which introgressed alleles are deleterious. We employed a similar parameter configuration to our simulations above: introgressed alleles are deleterious at 1,000 loci and cause a 20% fitness reduction in F1 hybrids. Neutral allele frequencies were tracked at 10,000 evenly-spaced loci. Under this configuration, for the recombination processes of humans and *D. melanogaster*, and for various genomic window sizes, we made the following observations (Fig. 3). (i) The positive correlation between recombination rate and introgressed ancestry builds up quickly, and then slowly dissipates over time (Fig. 3A). (ii) The correlation is stronger when taken across larger windows of the genome (Fig. 3A), as observed previously [e.g., Schumer et al. (2018)]. This is presumably because, in larger windows, the average introgressed fraction changes in a less stochastic fashion over time (and, in empirical settings, also because estimates of the introgressed fraction in larger windows are less noisy). (iii) The direct effect of recombination on the rate of purging of deleterious introgressed alleles is, at first, by far the more important mechanism in setting up the positive correlation between recombination rate and introgressed ancestry (Fig. 3B). Recombination’s effect in unlinking neutral from deleterious introgressed alleles gradually becomes more important, but for both humans and *Drosophila*, even 2,000 generations after admixture, it remains the minor mechanism (especially so for *Drosophila*).

**Figure 3:**
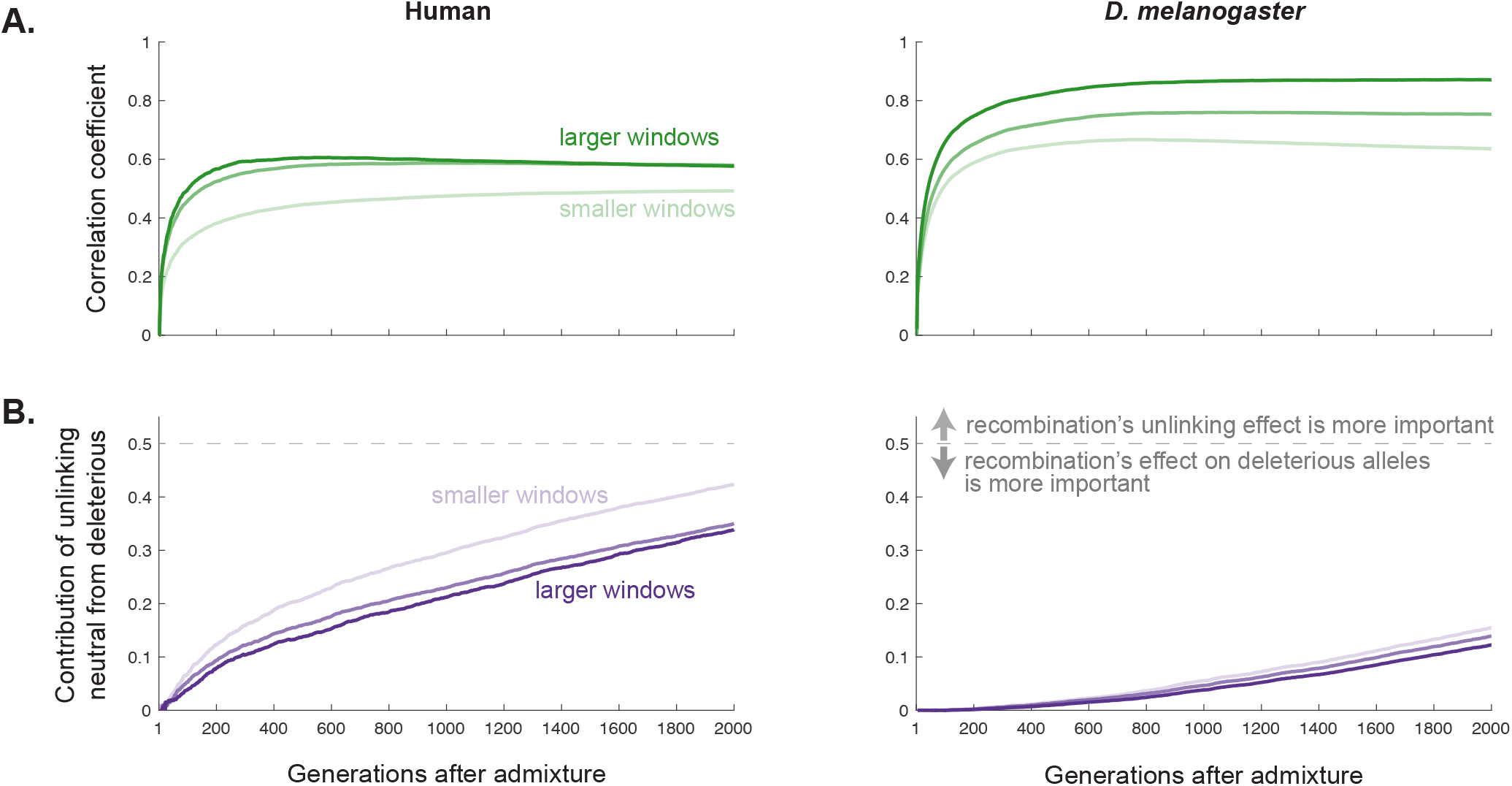
Genomic correlations between local recombination rate and introgressed ancestry. These simulations involve 10,000 evenly-spaced loci at which introgressed alleles are neutral, and 1,000 loci at which introgressed alleles are deleterious. The genome is divided into windows of size 10, 30, or 100 neutral loci, with windows contrained to lie on the same chromosome. Each generation, we calculate, across all windows, the correlation coefficient between the average introgressed fraction per window (calculated at neutral loci) and the average recombination rate between adjacent neutral loci in the window. We then use Eq. (17) to calculate the proportion of the correlation that is due to recombination’s effect in unlinking neutral from deleterious introgressed alleles. Profiles are averaged across 100 replicate simulations. **A.** The correlation between recombination rate and introgressed ancestry is set up quickly, and is stronger under the recombination process of *D. melanogaster* than of humans. **B.** For both recombination processes, recombination’s effect in unlinking neutral from deleterious introgressed alleles (thus decoupling their frequency trajectories) contributes little to the correlation between recombination rate and introgressed ancestry. Instead, for many generations after the admixture pulse, the correlation is driven by differences across the genome in the efficiency with which deleterious introgressed alleles are purged.

The reason for the relative unimportance of recombination’s unlinking effect is straightforward. Take a high-recombination case like humans, with ~20 chromosomes and ~1 crossover per chromosome per gamete. In the model as configured above, there would be ~50 deleterious loci per chromosome. Consider a neutral introgressed allele situated halfway between its two flanking deleterious alleles. For the neutral allele to dissociate from both its flanking deleterious alleles requires, first, a recombination event anywhere between the two deleterious alleles (rate ~1/50 → waiting time ~50 generations), which unlinks the neutral allele from one of the deleterious alleles, and then, subsequently, a recombination event between the neutral allele and the remaining linked deleterious allele (rate ~1/100 → additional waiting time ~100 generations). It therefore takes, on average, about 150 generations following the admixture pulse for the neutral allele to dissociate from both its flanking deleterious alleles (and even longer for neutral alleles that are closer to one flanking deleterious allele than the other).

As we have seen, by the time 150 generations have elapsed since admixture, most of the purging of introgressed ancestry has already occurred, and substantial ancestry differences have been set up between low-recombination and high-recombination regions. These differences across the genome, the imprint of which will persist for many generations, are therefore driven predominantly by the direct effect of recombination on the rate of purging of deleterious introgressed alleles.

The logic set out above is even more forceful in the case of low-recombination species like *Drosophila*: neutral alleles take even longer to dissociate from their flanking deleterious alleles, allowing even greater ancestry differences across the genome to be set up in the mean time by the direct effect of recombination on the rate of purging of deleterious alleles. This explains why, in our simulations of the *D. melanogaster* recombination process, 2,000 generations after the initial admixture pulse, the unlinking effect still accounts for less than 20% of the overall correlation between recombination rate and introgressed ancestry.

A corollary of the results above is that recombination’s role in determining the genome-wide rate of purging—and thus its role in generating species differences in the retention of introgressed DNA—is driven largely by recombination’s impact on the rate of purging of deleterious introgressed alleles, rather than its effect in unlinking neutral from deleterious introgressed alleles. Even though most introgressed DNA is expected to be neutral, the persistence of the neutral alleles’ initial linkage to deleterious alleles causes them to be purged genome-wide at a rate that is, for many generations, almost identical to the rate of purging of the deleterious alleles themselves (Fig. S5).

## 5 Discussion

Recent genomic evidence has shown that, following admixture, introgressed alleles tend to be deleterious at many loci throughout the recipient species’ genome. Here, we have studied the influence of the aggregate recombination process on the efficiency with which selection purges introgressed DNA genome-wide. We have shown that species (and genomic regions) with low aggregate recombination rates—as quantified by 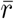 and analogous metrics—purge introgressed DNA more rapidly and more profoundly than species (and genomic regions) with high aggregate recombination rates. For most species, these effects are driven predominantly by recombination’s effect on the rate of purging of deleterious introgressed alleles, rather than its effect in unlinking neutral introgressed alleles from their deleterious counterparts.

### 5.1 Empirical predictions

The simplest prediction emerging from our analysis is that species with fewer chromosomes should exhibit a weaker genomic signal of historic introgression. This is because (i) species differences in the retention of introgressed DNA are typically set up in the first few generations after hybridization, when most purging of introgressed DNA occurs (Fig. 1); and (ii) the rate of purging in these first few generations is governed by the aggregate recombination rate, which is dominated by the effect of independent assortment of chromosomes and thus by the number of chromosomes (Veller et al. 2019).

Directly testing this prediction is not yet possible, because quantitative estimates of genome-wide introgressed fractions have been obtained for only a handful of species. However, with the rapid accumulation of sequence data from many taxa and the development of multiple complementary methods to identify introgressed tracts within sequence data (Dagilis et al. 2021), we are confident that our prediction will soon be testable. A particularly promising clade in this regard is *Drosophila*, because, with its small baseline karyotype, the variation in chromosome number observed in the genus (Bracewell et al. 2019) corresponds to substantial variation in aggregate recombination rate. For example, *D. melanogaster* has only two major autosomes, the independent assortment of which contributes ~0.25 to 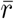. *D. subobscura*, in contrast, has four major autosomes, the independent assortment of which contributes ~0.38 to 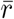 [this calculation makes use of chromosome lengths reported by Bracewell et al. (2019)]. This karyotypic advantage, together with substantial progress in the analysis of introgression across the genus [e.g., Suvorov et al. (2021)], suggests that *Drosophila* will likely be among the most informative clades in which to test the prediction that chromosome number correlates positively with introgressed ancestry.

In the mean time, we can make use of one kind of data that has been extensively collected in cases of hybridization: the geographic width of hybrid zones. Our general prediction that species with fewer chromosomes should experience more efficient selection against hybrid ancestry, and therefore more rapid purging of introgressed DNA, carries the corollary that hybrid zones should be narrower between species with fewer chromosomes. McEntee et al. (2020) collated hybrid zone widths for 131 animal species pairs, together with estimates of genetic divergence and dispersal distance. They used these data to show that dispersal distance and genetic divergence alone can explain much of the variation in cline widths. We obtained karyotype data for 54 of the 131 species pairs (details in SI File S1). Rerunning the regression of McEntee et al. (2020) on this reduced sample of 54 species pairs (Table 1), and including as additional independent variables several estimates of aggregate recombination derived from the karyotypes, we found that inter-chromosomal 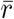 (the portion due to independent assortment of chromosomes) was positively associated with cline width, as predicted. This result was not statistically significant at conventional levels (*p* = 0.13), which is perhaps unsurprising given the small sample size. Encouragingly, the 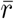-based proxy for aggregate recombination was a stronger predictor of cline width than alternative proxies such as chromosome number and the effective number of chromosomes [the number of evenly sized chromosomes required to match the observed inter-chromosomal 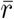 (Szymura and Barton 1986; Waples et al. 2021)]. Interestingly, the coefficient on a binary variable for when the hybridizing species pair have unequal chromosome numbers was significantly negative (*p* = 0.007), consistent with selection against karyotype heterozygotes in hybrid zones.

**Table 1:**
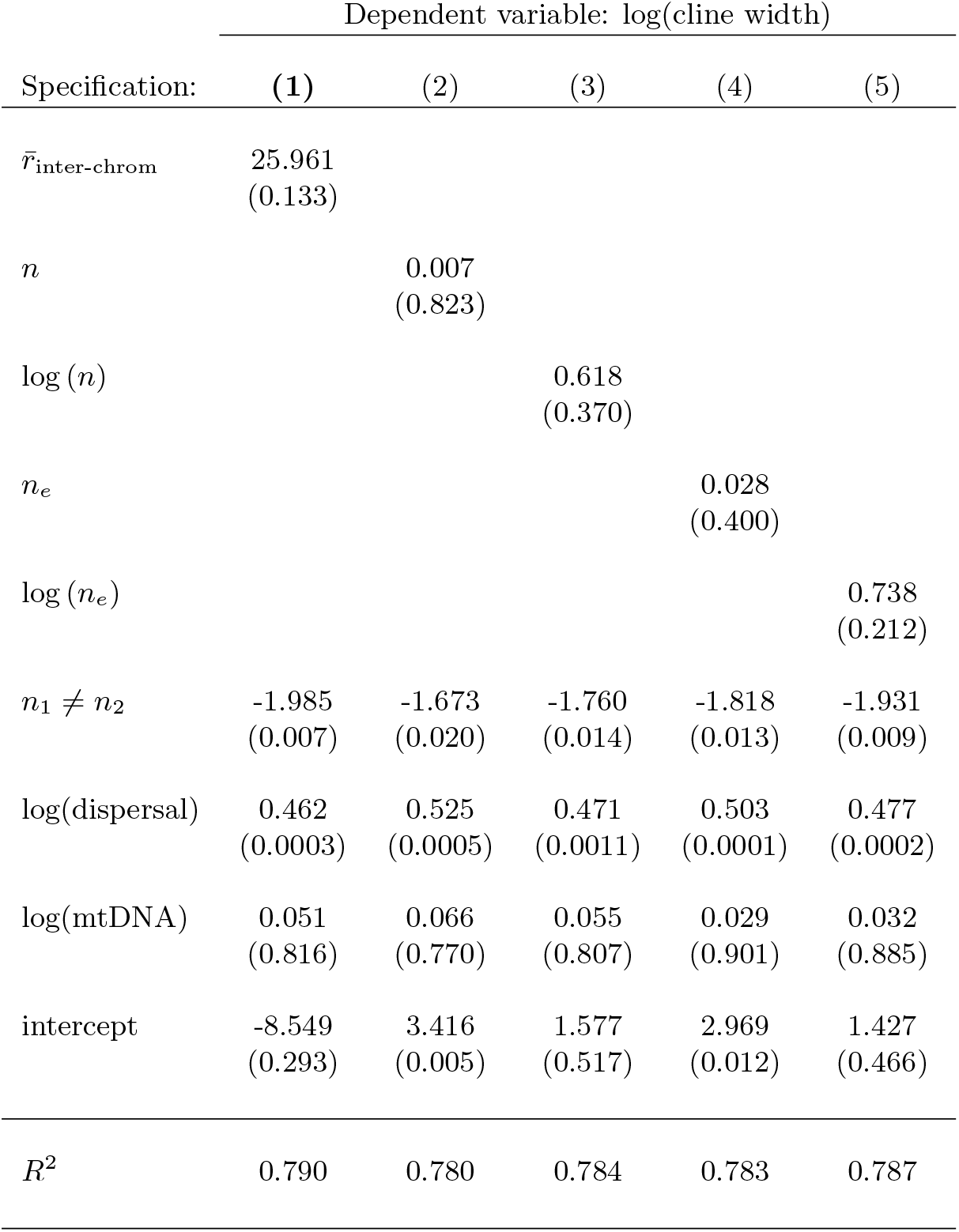
Effect of chromosome number on cline width across 54 hybridizing pairs of vertebrate species. Coefficient estimates from least squares regression of cline width (km) on dispersal distance (km), genetic divergence (distance between mitochondrial sequences), several metrics of the amount of genetic shuffling caused by independent assortment of chromosomes, and a binary variable for whether the hybridizing pair have unequal chromosome numbers. *p*-values in brackets underneath coefficient estimates. Data for cline widths, dispersal distances, and genetic divergence estimates are from McEntee et al. (2020). Karyotype data are listed in SI File S1. **Key:** 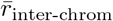: Contribution to 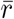 from independent assortment of chromosomes in the homogametic sex, taking into account the number and relative sizes of the chromosomes. When karyograms were available for both species, the average value was used. *n*: Average number of chromosomes for the species pair. *n_e_*: Effective number of chromosomes: The number of evenly sized chromosomes required to generate an interchromosomal value of 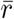 equal to the species’ true value, calculated as 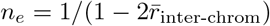. *n*_1_ ≠ *n*_2_: Indicator variable that takes the value 1 if species 1’s chromosome number, *n*_1_, is different to species 2’s chromosome number, *n*_2_, and takes the value 0 otherwise.

| Specification: | Dependent variable: log(cline width) |  |  |  |  |
| --- | --- | --- | --- | --- | --- |
|  | (1) | (2) | (3) | (4) | (5) |
| $\bar{r}_{\text{inter-chrom}}$ | 25.961<br>(0.133) | | | | |
| $n$ | | 0.007<br>(0.823) | | | |
| $\log(n)$ | | | 0.618<br>(0.370) | | |
| $n_e$ | | | | 0.028<br>(0.400) | |
| $\log(n_e)$ | | | | | 0.738<br>(0.212) |
| $n_1 \neq n_2$ | -1.985<br>(0.007) | -1.673<br>(0.020) | -1.760<br>(0.014) | -1.818<br>(0.013) | -1.931<br>(0.009) |
| $\log(\text{dispersal})$ | 0.462<br>(0.0003) | 0.525<br>(0.0005) | 0.471<br>(0.0011) | 0.503<br>(0.0001) | 0.477<br>(0.0002) |
| $\log(\text{mtDNA})$ | 0.051<br>(0.816) | 0.066<br>(0.770) | 0.055<br>(0.807) | 0.029<br>(0.901) | 0.032<br>(0.885) |
| intercept | -8.549<br>(0.293) | 3.416<br>(0.005) | 1.577<br>(0.517) | 2.969<br>(0.012) | 1.427<br>(0.466) |
| $R^2$ | 0.790 | 0.780 | 0.784 | 0.783 | 0.787 |

### 5.2 Introgression selects for lower recombination

We have shown that aggregate recombination affects the rate at which deleterious introgressed DNA is purged. Since we know hybridization and subsequent genetic introgression to be common, this raises the converse question: does introgression select for modification of the recombination process?

Introgression’s effect on modifiers of local recombination rates is straightforward. A modifier allele in the recipient species that reduces its local recombination rate prevents deleterious introgressed alleles from recombining onto its background, and is thus favored by selection. For example, a segregating inversion keeps together a haplotype of non-introgressed alleles, and is therefore favored over the alternative haplotype whose orientation is the same as that in the donor species and which therefore admits deleterious introgressed alleles by recombination (Kirkpatrick and Barton 2006).

Our results also point to how selection acts on global modifiers of the recombination process in the face of deleterious introgression. A modifier allele that reduces the aggregate recombination rate (i.e., 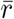 and analogous metrics) increases the variance among its descendants in how much introgressed DNA they carry (Veller et al. 2020). This allows selection to purge introgressed DNA more efficiently among descendants of the modifier allele, causing the modifier allele to end up in fitter genotypes and thus to be positively selected. This logic is similar, but the conclusion opposite, to the usual case where a global modifier that increases the recombination rate is favored by selection because it increases fitness variance among its descendants (Barton 1995; Burt 2000; Barton and Otto 2005). The conclusions are opposite because, in the usual case, the interaction of selection and random drift generates, on average, negative linkage disequilibria between deleterious alleles (Barton and Otto 2005), whereas in our case, the deleterious alleles are introgressed into the recipient population in perfect positive linkage disequilibrium.

Therefore, introgression generates selection on both local and global modifiers to reduce the recombination rate. Local modifiers of recombination include structural rearrangements (Kirkpatrick 2010), alterations to the binding sites of recombination-specifying proteins (Paigen and Petkov 2018; Grey et al. 2018), and mutations that affect local chromatin structure in meiotic prophase [e.g., Stack et al. (2017)]. The simplest global modification of the aggregate recombination rate is a change in chromosome number (Veller et al. 2019), but introgression is not expected to select for reduced chromosome number owing to fertility problems in karyotype-heterozygous hybrids (White 1978). Therefore, introgression is expected to select for global modification of the recombination process only via modifiers of the number and spatial arrangement of crossovers. Our expanding knowledge of the molecular biology of meiosis and recombination [reviewed in Hunter (2015); Zickler and Kleckner (2015)] suggests such modifiers to be very common: They include mutations to key meiosis proteins, such as those that determine the lengths of chromosome axes in meiotic prophase [e.g., Novak et al. (2008); Hong et al. (2019)], those that control the interference process along chromosome axes [e.g., Zhang et al. (2014)], and those that globally specify recombination hotspots (Paigen and Petkov 2018; Grey et al. 2018).

Frequent introgression is therefore expected to shape genetic variation in these factors towards reducing both local and global recombination. In this way, selection for reduced recombination acts as an indirect form of reinforcement, causing post-zygotic selection against introgressed DNA to be more efficient, and thus strengthening the barrier to gene flow between species.

## Acknowledgements

We are grateful to Graham Coop for many helpful discussions and suggestions on the content and framing of this manuscript, and to Erin Calfee, Jeff Groh, Jim Mallet, Molly Schumer, and Sivan Yair for helpful comments. CV is supported by a Branco Weiss fellowship. NBE is supported by a G. Evelyn Hutchinson Environmental Postdoctoral Fellowship. PM is supported by a Center for Population Biology postdoctoral fellowship and an NSF postdoctoral fellowship. This work was supported in part by the National Institute of General Medical Sciences of the National Institutes of Health (grants NIH R01 GM108779 and R35 GM136290 to G. Coop).

## S1 A breeder’s equation for the rate of purging of introgressed DNA

Let the random variable *Z_t_* be the fraction of introgressed DNA carried by a generation-*t* zygote, and let the random variable *G_t_* be the fraction of introgressed DNA in a successful gamete from generation *t* (i.e., after selection has acted). The relative fitness of a zygote with introgressed fraction *Z* is 1 − *ZS*. Suppose that the probability density function for *Z_t_* is *p*(*x*). Then the probability density function for *G_t_* is

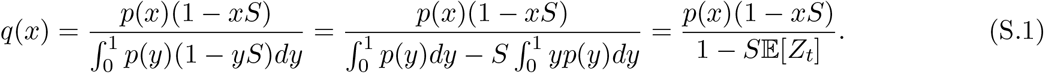

From this,

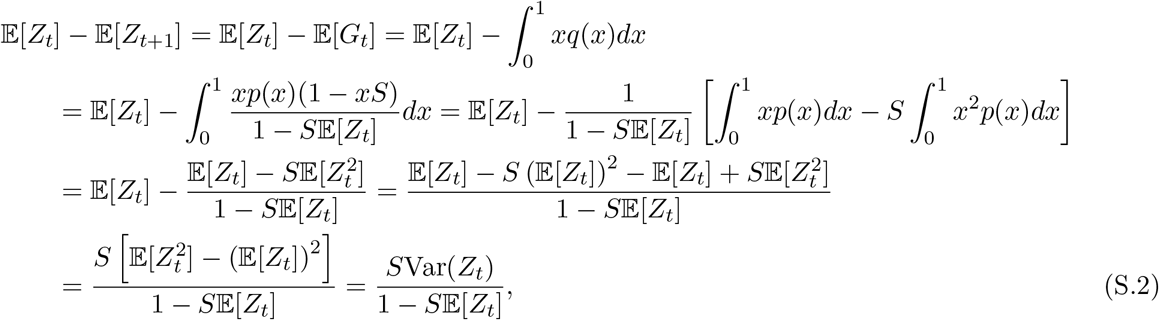

which is Eq. (1) in the Main Text.

## S2 A model to distinguish the contributions of block number variance and block length variance to the overall rate of purging

We define blocks of introgressed DNA as sets of introgressed alleles with identical inheritance pedigrees going back to the admixture pulse. That is, they need not be contiguous stretches of the genome (and most blocks will not in fact be contiguous in the first few generations after admixture). We can distinguish two sources of variance across individuals in how much introgressed DNA they carry: (i) variance in the number of blocks of introgressed DNA carried; (ii) variance in the length of blocks. We are interested in the effect of recombination on these two sources of variance, and the relative importance of these two sources for the overall ancestry variance (and thus the rate of purging of introgressed DNA).

Here, two timescales must be distinguished. In the early generations after admixture, the number of blocks that an individual carries depends largely on its pedigree, and not on the particular patterns of recombination within that pedigree. Thus, for example, a generation-2 individual will almost certainly have inherited zero, one, or two blocks of introgressed DNA if, respectively, zero, one, or two of its (generation-1) parents were F1 hybrids. Since the blocks that it inherits (if any) are very large, it too will transmit zero, one, or two blocks to its offspring, regardless of the species’ particular recombination process. Thus, in these very early generations after admixture, recombination affects ancestry variance across individuals only by affecting block length variance, and not by affecting block number variance.

In later generations, however, introgressed blocks become sufficiently mixed among the population that they can be assumed to have been inherited approximately independently from one another. Then recombination affects variance both in the number of blocks carried by different individuals and in the lengths of these blocks, in a way that we shall now characterize formally.

In the population of size *N*, in a given generation, suppose that the overall proportion of introgressed ancestry is *x* (which we assume to be small). The introgressed DNA is contained in a total of *n* blocks of average length 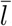, measured as a fraction of total diploid genome length, so that 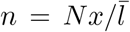. Since blocks can approximately be assumed to have been inherited independently, the probability that an individual gets a particular block is 1*/N*, and so the number of blocks that a given individual gisetbsi,n*B*om, ially distributed with p rameters *n* and *p* = 1*/N*. Therefore, 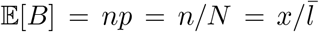 and 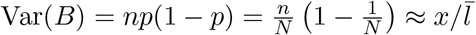 when *N* is large.

Let the random variable *Z* be the fraction of an individual’s genome that is introgressed (so that 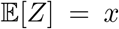). Then 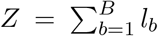, where *l_b_* is the length of the *b*-th block assigned to the individual. 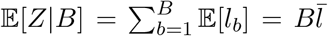 and, because the *l_b_* are independent, 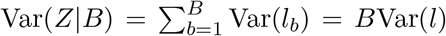. From the law of total variance,

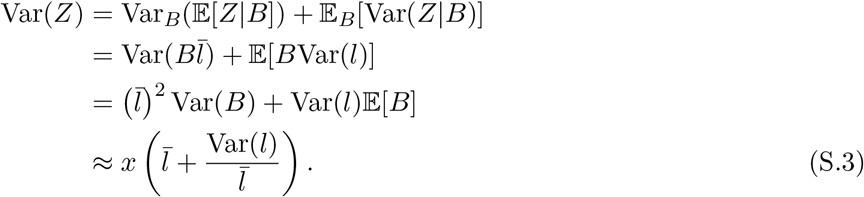

The first term in Eq. (S.3) is the contribution to variance in *Z* from variance in the number of blocks carried by different individuals, while the second term is the contribution from variance in the lengths of different blocks.

From Eq. (S.2), the proportion of introgressed ancestry that is purged from one generation to the next is

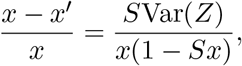

where *S* is the fitness reduction of individuals with 100% introgressed ancestry. Substituting in the variance calculation above,

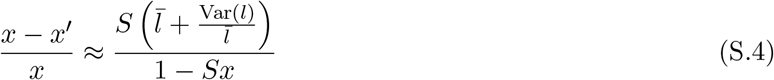

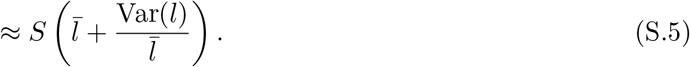

So the rate of purging depends only on the average and variance of the block lengths. Eq. (S.5) can be written in simpler form:

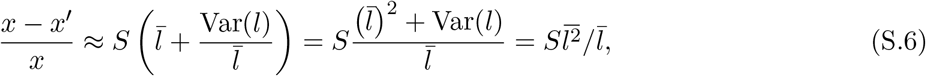

where 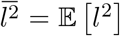 is the uncentered second moment of the distribution of block lengths. Fig. S1A shows the numerical accuracy of Eq. (S.4) for *Drosophila*’s recombination process, in the case where all F1s are hybrids (so that the level of [recombination-independent] block number variance in the early generations is minimal).

In the early generations after hybridization, the aggregate recombination process does not generate substantial block number variance, for reasons discussed above, but it can generate substantial block length variation. For example, if crossovers tend to be situated near the ends of chromosomes, or if the chromosomes themselves vary greatly in size, then the blocks into which recombination and segregation break introgressed ancestry will tend to be more variable in size. In contrast, eventually recombination will have dissociated all introgressed alleles from one another, so that every block is one locus long (*l_∞_* = 1*/* [2*L*]). Then 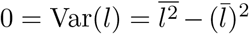 so that 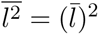, and the rate of purging is 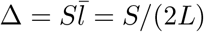, the asymptotic value observed in our simulations (e.g., Fig. 1B). [This asymptotic rate will be reached as long as the population is not so small that genetic drift dominates selection on individual alleles (*Ns* ≪ 1).] Therefore, eventually, all of the ancestry variance across individuals is due to (single-locus) block number variance. This is in contrast to the early generations, where block length variance contributes substantially to overall ancestry variance.

We can compare the contributions of these two sources of variance formally using the decomposition in Eq. (S.3). The contribution of block length variance relative to block number variance is

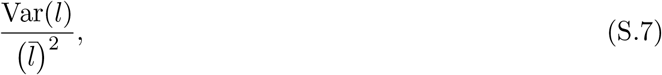

which is simply the squared coefficient of variation of block lengths. Fig. S1B plots this quantity over time in the case where all F1s are hybrids, and, together with the arguments above, reveals that block length variance is particularly important for the purging of introgressed DNA in the early generations after hybridization. Over time, it becomes less important, with block number variance eventually becoming the only source of ancestry variance across individuals.

The analysis above permits the following interpretation of the genomic scales of recombination that are most important for the rate of purging of introgressed ancestry at different timescales after admixture.

**Figure S1:**
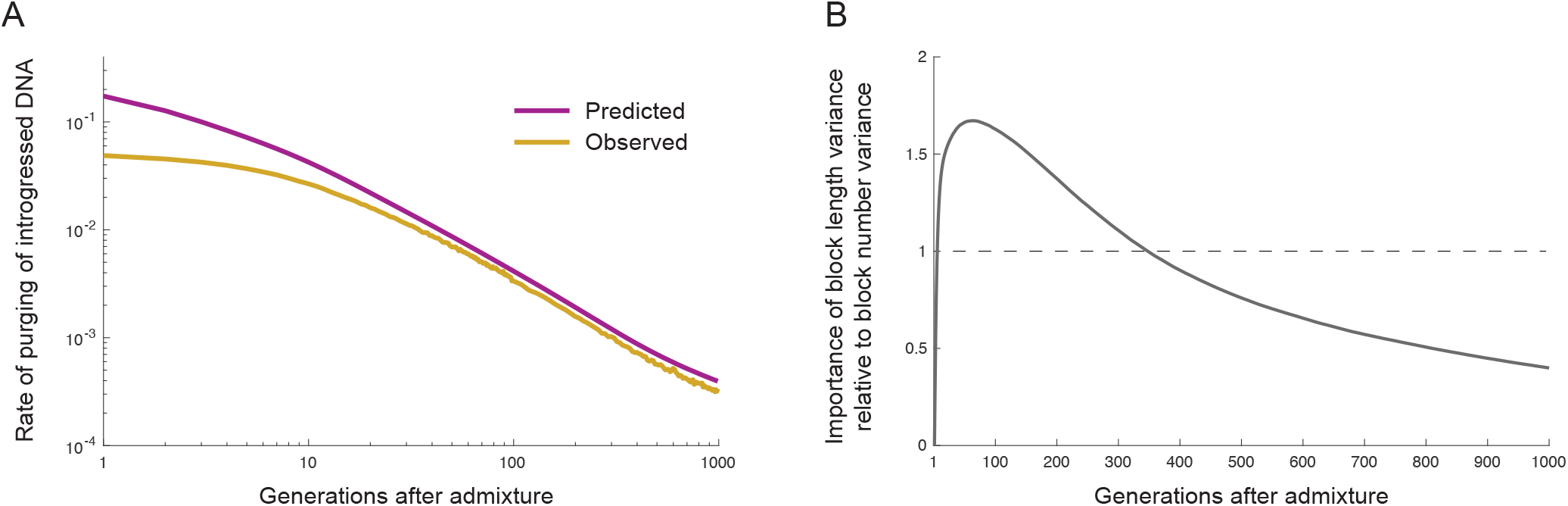
The rate of purging of introgressed DNA is determined by the distribution of introgressed block lengths. The recombination process is that of *D. melanogaster*, and we assume that all F1s are hybrids to eliminate recombination-independent variance in block number in the early generations after admixture. Trajectories are averaged across 1,000 replicate simulations. For computational reasons, the population size is *N* = 10,000, rather than the value of *N* = 100,000 used for simulations in the Main Text. **A**. Observed rate of purging vs. the prediction of Eq. (S.4). Eq. (S.4) overestimates the rate of purging in the generations immediately after hybridization, because the sampling process on which it is based involves block number variance across individuals when in fact there is little/none in these early generations. Eq. (S.4) becomes more accurate in later generations as blocks become more and more randomly mixed among the population. **B**. Relative importance of the two sources of variance in introgressed ancestry, calculated using Eq. (S.7). Block length variance is important in the early generations, implicating the aggregate recombination process in the purging of introgressed DNA, while block number variance is more important in the later generations, implicating fine-scale recombination rates. Although the figure does not display this, block length variance is in fact substantially more important than block number variance in the earliest generations after hybridization, for reasons explained in the text. The discrepancy arises because, again, the assumptions of the model used to derive Eq. (S.7) do not hold in the earliest generations after admixture.

In the first few generations after hybridization, recombination affects the rate at which introgressed DNA is purged because it chops the initial linkage blocks into smaller blocks of variable size. This implicates the aggregate recombination process—in particular, heterogeneity in chromosome size and the spatial distribution of crossovers—in the early purging of introgressed DNA [cf. Eq. (10)]. In later generations, recombination affects the rate at which introgressed DNA is purged primarily because it affects block number variance, which is proportional to average block length [Eq. (S.3)]. Therefore, the key effect of recombination in later generations is simply to chop blocks up into more blocks: when there is crossing over anywhere within a block, the recombinant block that segregates to a given gamete has average size one half that of the original block, no matter the spatial position of the crossover within the block. This implicates the fine-scale recombination rate (cM/Mb) in later-generation purging.

In summary, the aggregate recombination process (as quantified by 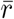 and analogous metrics) is most important in the early generations after hybridization (when most purging occurs), while the fine-scale recombination process is most important in the later generations. This mirrors the implications we drew from Eq. (16) in the Main Text for the case where selection against introgressed ancestry is weak and the initial introgressed fraction is small.

**Figure S2:**
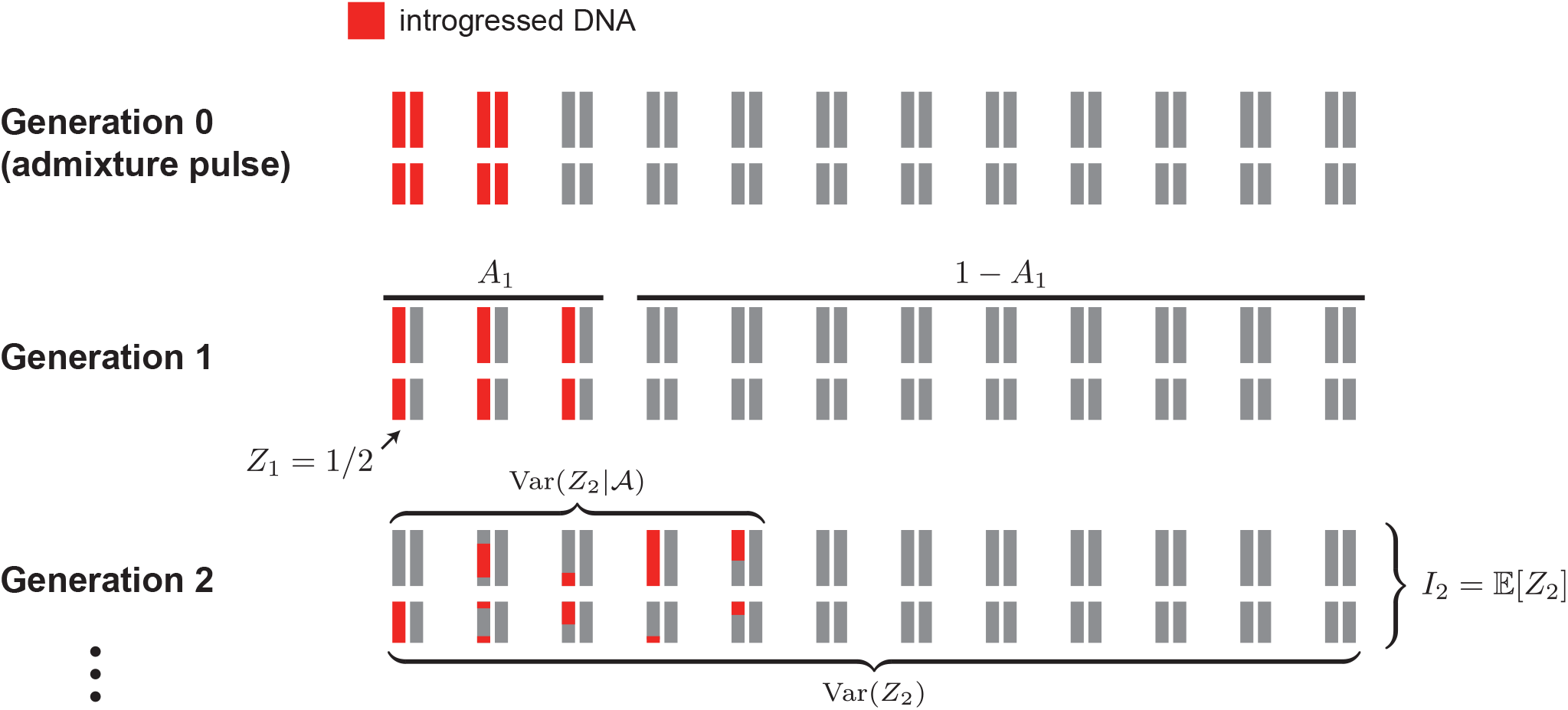
Graphical illustration of the variables used in the Main Text calculations.

**Figure S3:**
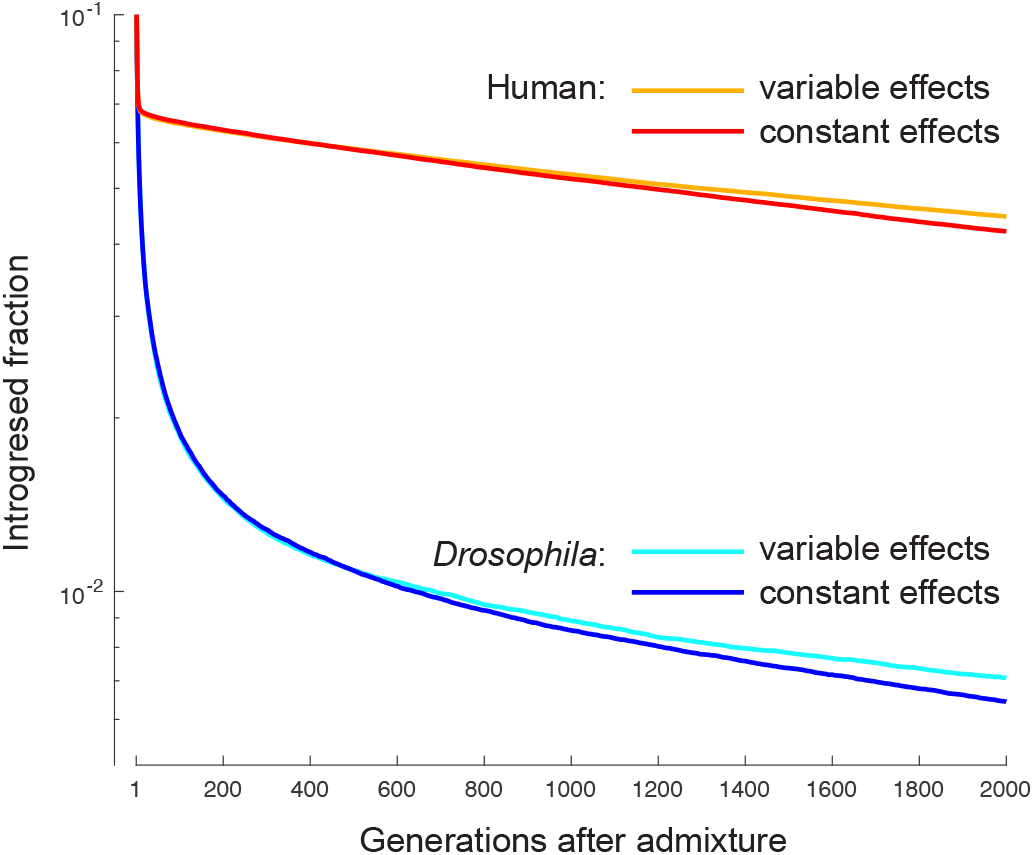
Purging of deleterious introgressed DNA when the deleterious alleles have constant effect sizes vs. variable effect sizes. In the variable case, the effect size of the introgressed allele at each locus is drawn independently from an exponential distribution whose mean is equal to the effect size in the constant-effect-size case (2 × 10^−4^). The rate of purging is similar between the two cases when block lengths are still large, because the mean allelic effect size is then most important. Later, the rate becomes slower in the variable effect size case, because large-effect alleles have been preferentially purged so that the average effect size has declined below its initial value (which always remains the average value in the constant effect size case). Nonetheless, the impact of allowing variable effect sizes is small. Trajectories are averaged across 100 replicate simulations.

**Figure S4:**
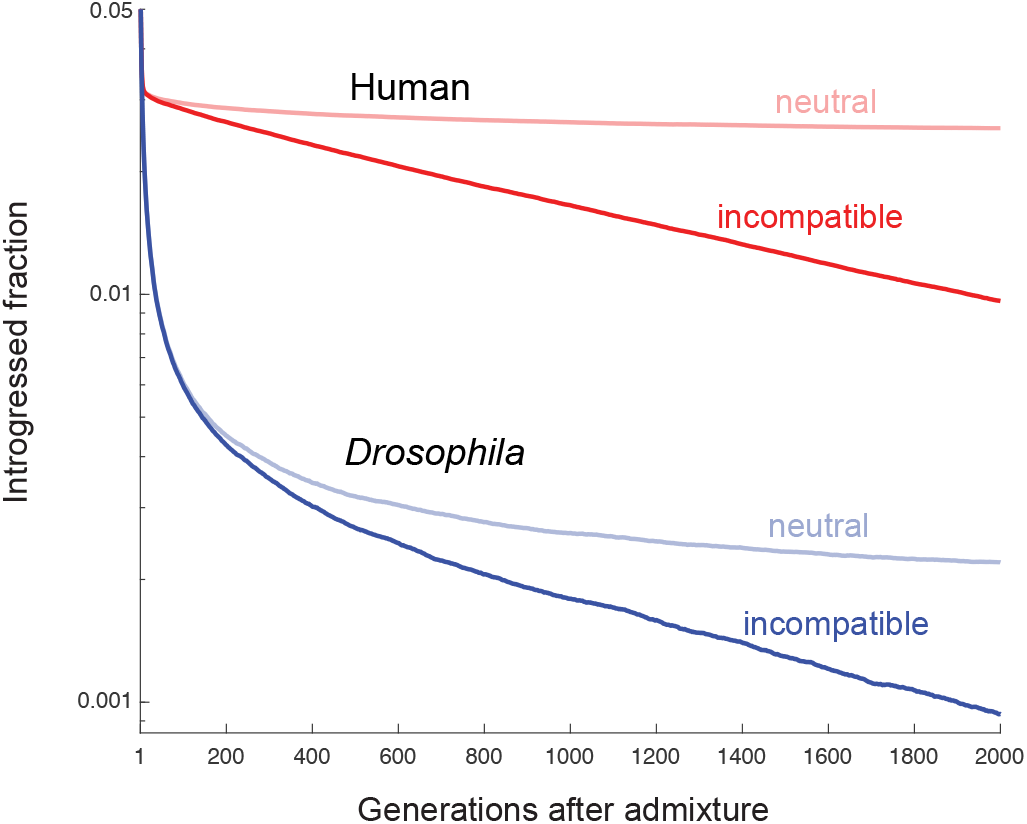
Purging of introgressed alleles when some of them are incompatible with recipient-species alleles. In the simulations here, there are 500 Dobzhansky-Muller incompatibilities (DMIs), each involving a distinct, randomly chosen locus pair among 1,000 loci spaced evenly along the physical map of the genome. We assumed intermediate epistatic dominance for all DMIs: If *A* and *B* are the incompatible alleles at a locus pair, with *a* and *b* the alternative alleles, then the genotype *AABB* suffers a relative fitnes reduction 1 − *s*, the genotypes *AaBB* and *AABb* a relative fitness reduction 1 − 3*s/*4, and the genotype *AaBb* 1 − *s/*2. The value of *s* was chosen such that F1s suffer an overall fitness reduction of 20%, as in our other simulations. Solid lines show the average introgressed fraction at the 500 loci where the introgressed alleles are incompatible with recipient-species alleles elsewhere. Neutral ancestry fractions (faded lines) were tracked at 10,000 evenly-spaced loci. Trajectories are averaged across 100 replicate simulations.

**Figure S5:**
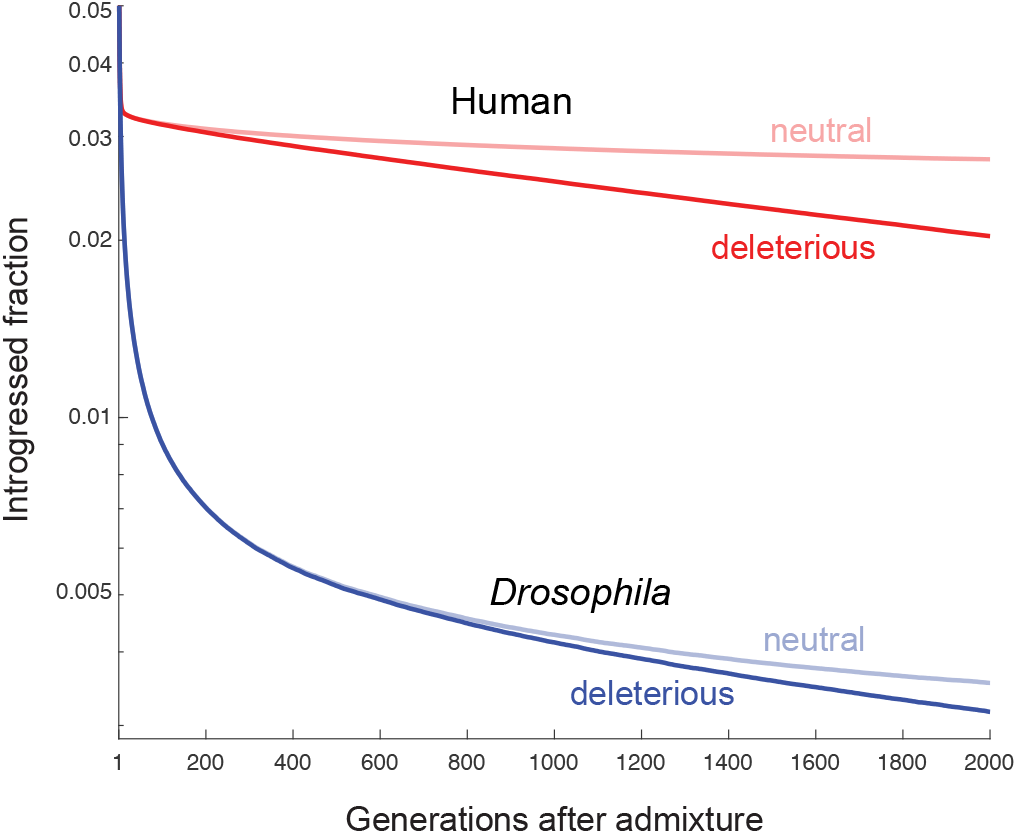
When there are sufficiently many loci at which the introgressed alleles are deleterious (here, 1,000), the frequency dynamics of deleterious and neutral introgressed ancestry are similar. Under the human recombination process, the average trajectories of deleterious and neutral introgressed ancestry begin to diverge appreciably from one another after ~200 generations; for *D. melanogaster*, this takes ~800 generations. In these simulations, introgressed alleles are deleterious at 1,000 evenly spaced loci, and neutral ancestry is measured at 10,000 evenly spaced loci. Trajectories are averaged across 100 replicate simulations.

**Figure S6:**
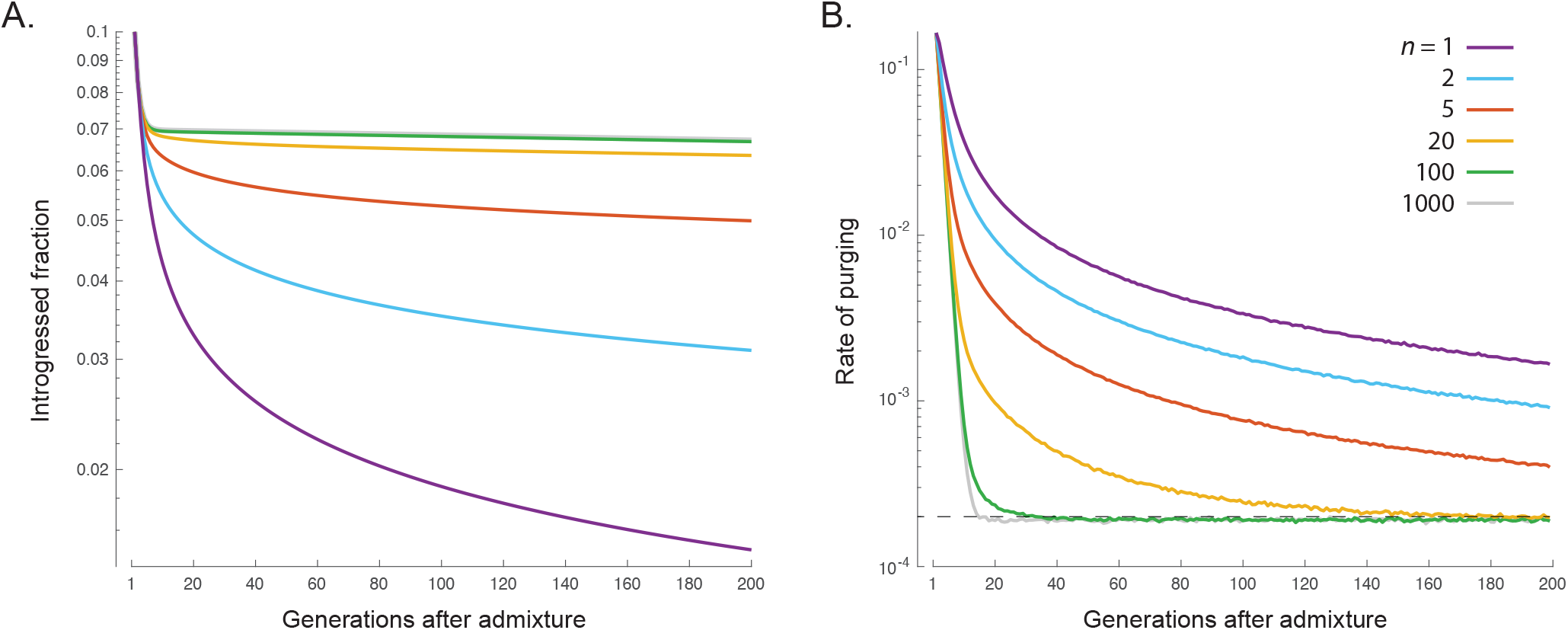
The effect of chromosome number on the purging of introgressed DNA. The model simulated here involves *n* chromosomes of equal size, with 1,000 loci distributed equally among the chromosomes and spaced evenly along them. There is, on average, one crossover per chromosome per gamete, with crossover positions uniformly distributed along the chromosome. When *n* is larger, the purging of introgressed DNA is substantially slowed, most obviously in the short run (owing to higher aggregate recombination—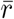 and analogs—caused largely by independent assortment of a greater number chromosomes) but also in the medium run (owing to a higher average fine-scale recombination rate, caused by a greater number of crossovers). Note, however, that the rate of purging in each case will eventually converge to the average allelic effect (dotted line in **B**), which is *s* = 2 × 10^−4^ in the specification considered here. Trajectories are averaged across 100 replicate simulations.

**Figure S7:**
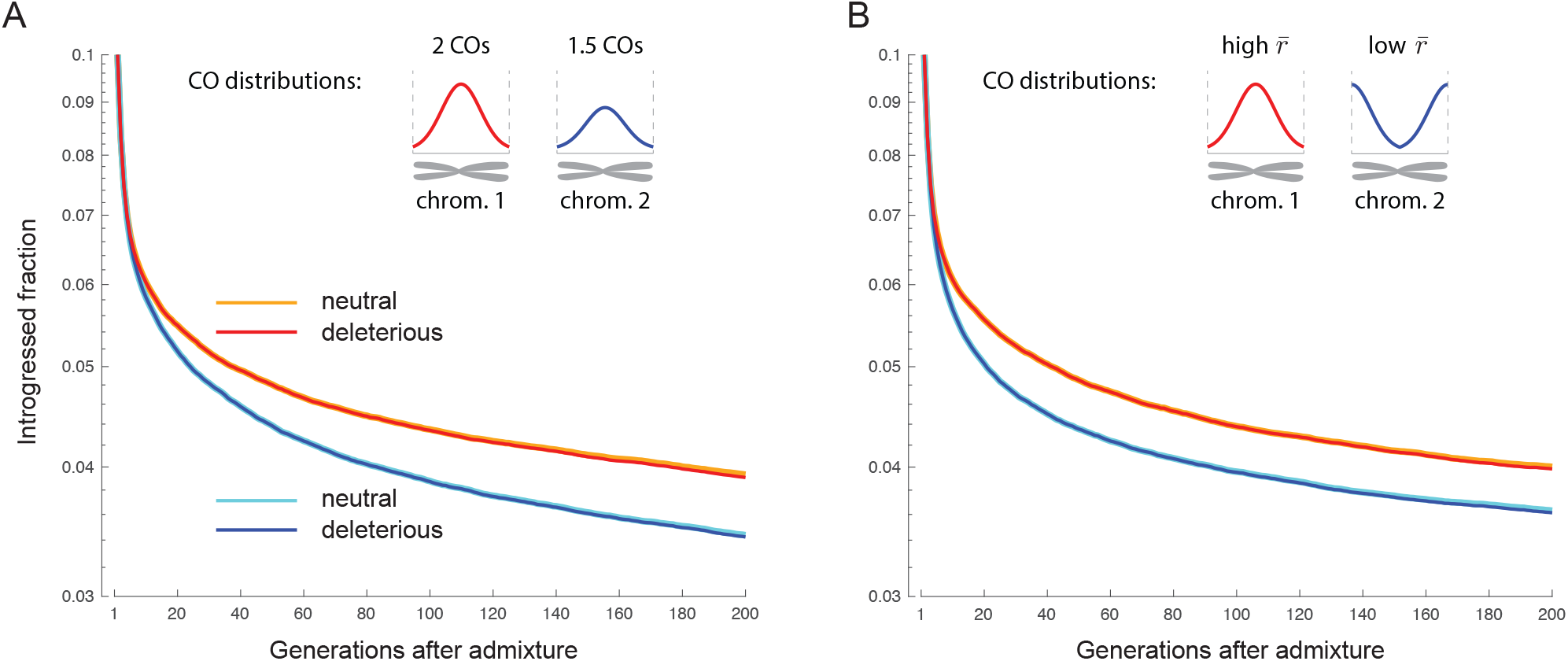
Genomic regions with low aggregate recombination purge more introgressed DNA. In these simulations, the genome comprises 10,000 evenly spaced loci spread equally across 2 chromosomes. Introgressed alleles are deleterious at 1,000 evenly spaced loci. **A**. Chromosomes 2 experiences, on average, 25% fewer crossovers than chromosome 1. The spatial distribution of crossovers is the same for the two chromosomes. The smaller number of crossovers causes both the aggregate and fine-scale recombination rates of chromosome 2 to be lower, and so the rate of purging of introgressed DNA is higher for chromosome 2, both in the early generations after hybridization, and later on. **B**. Chromosomes 1 and 2 experience the same average number of crossovers per meiosis (2 crossovers), but these crossovers tend to be more terminally localized on chromosome 2, causing chromosome 2 to have a lower aggregate rate of recombination (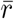 and analogs) than chromosome 1. Because of this, the rate of purging of introgressed DNA is initially higher on chromosome 2 than on chromosome 1. Because chromosome 1 and chromosome 2 have equal average fine-scale recombination rates, their rates of purging become similar in later generations. Nonetheless, the early rate difference leads to chromosome 1 ultimately retaining substantially more introgressed ancestry than chromosome 2. In both **A** and **B**, the average frequency trajectories of neutral introgressed ancestry closely resemble the trajectories of deleterious introgressed ancestry, for reasons explained in the Main Text. Therefore, the effect of recombination on differences in neutral introgressed ancestry across the chromosomes is driven almost entirely by the effect of recombination on the purging of deleterious introgressed alleles, rather than recombination’s effect in unlinking neutral introgressed alleles from their deleterious flanking alleles—see Fig. S8. Trajectories are averaged across 100 replicate simulations.

**Figure S8:**
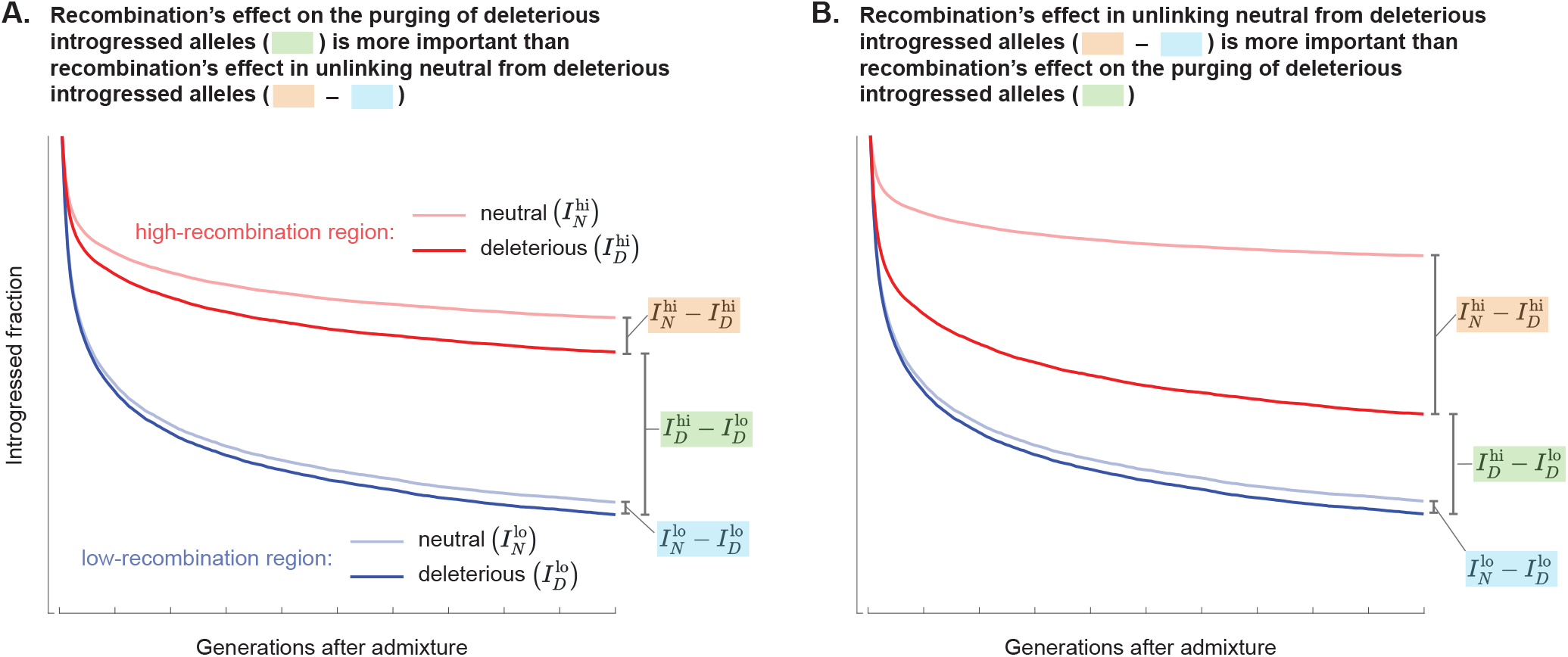
Graphical representation of the decomposition of the overall genomic correlation between recombination rate and introgressed ancestry [Eq. (17)]. In regions of low recombination, deleterious introgressed alleles are purged more rapidly, so the bold blue curves in **A** and **B** lie below the bold red curves, by an amount that quantifies this ‘channel’ by which the correlation between recombination and introgressed ancestry is set up. Also in regions of low recombination, neutral introgressed alleles remain in linkage with deleterious introgressed alleles for longer, so that the faded blue curves lie closer to the bold blue curves than the faded red curves lie to the bold red curves—the disparity between these two gaps quantifies the ‘unlinking’ channel by which the correlation between recombination and introgressed ancestry is set up.

## Notes

### Competing Interest Statement

The authors have declared no competing interest.

### Summary of Updates

An earlier version of this manuscript had two parts: (1) Calculations of the variance of genetic relatedness between individuals with particular pedigree relationships, taking into account the randomness of recombination and segregation in their pedigree. (2) An investigation of the rate of purging of introgressed DNA following admixture, based in part on results from part (1). Part (1) has since been published as Veller et al. (2020) Genetics, 216(4), 985-994. The present manuscript has been reconfigured to focus on part (2).

